# Genomic adaptations of novel halotolerant bacteria from extreme North Greenland

**DOI:** 10.1101/2025.06.13.657406

**Authors:** Miguel Ángel Salinas García, Anders Priemé

**Affiliations:** Centre for Exolife Sciences (CELS), Niels Bohr Institute, University of Copenhagen, Jagtvej 155A, DK-2200, Copenhagen, Denmark; Center for Volatile Interactions (VOLT), Department of Biology, University of Copenhagen, Universitetsparken 15, DK-2100 Copenhagen, Denmark; Section of Microbiology, Department of Biology University of Copenhagen, Universitetsparken 15, DK-2100 Copenhagen, Denmark

**Author notes:** **Corresponding author**: Anders Priemé. Mailing address: Center for Volatile Interactions (VOLT), Department of Biology, University of Copenhagen, Universitetsparken 15, DK-2100 Copenhagen, Denmark.

**Keywords:** extremophile, Greenland, biosynthetic gene cluster, halotolerant, genome, novel strains

## Abstract

Peary Land, northern-most Greenland, is a cold desert region whose soil microbiome remains underexplored. This is the first study to explore the halotolerant soil microbiome of this region with culture-dependent methods. Forty-nine taxonomically diverse bacterial isolates were obtained from dry biological soil crust, surface soil and permafrost soil. Nine isolates were selected for whole genome sequencing; these genomes showed adaptations to their cold, saline native environment, such as cold shock proteins and a large number of osmoprotectant transporters. The genomes also contained several biosynthetic gene clusters, including a large variety of clusters involved in the production of antimicrobial compounds. Furthermore, most of the sequenced genomes showed low digital DNA-DNA hybridisation to their closest relatives, suggesting that they represent novel species. Overall, this study demonstrates the unexplored potential of High Arctic deserts as a source of novel halotolerant bacteria and their secondary metabolites.

## INTRODUCTION

Peary Land in extreme North Greenland is the northern-most landmass of our planet. The interior of this remote and uninhabited peninsula is characterized by very sparse vegetation, very low precipitation and very low temperatures [1]. Due to the low precipitation, most of Peary Land is ice-free and vegetation is mainly found along the moist northern coast bordering the icy Arctic Ocean and in small depressions in the landscape fed by water from melting snowpack. In addition, Peary Land is home to one of the oldest permafrost formations in the world, located in Kap København and around 2 million years old [2]. Other notable locations include Cape Morris Jessup, the most northern point of mainland Greenland, and Citronen Fjord, which contains rich deposits of zinc and lead and is being considered for exploitation (Ironbark Zinc ltd.).

Due to the low precipitation and sparse vegetation, the soils in Peary Land are generally thin, alkaline and salt-rich, and poor in carbon, nitrogen and phosphorus [3]. Surface soils of the High Arctic desert of Peary Land tend to harden and crack due to lack of water and evaporation, creating the typical patterns found in deserts [4]. In contrast to warm or hot deserts, UV radiation is relatively low, with noon UV index averages of 2-3 in July [5, 6].

Biological soil crusts are common in hot and cold deserts and have been defined as communities of organisms dwelling on the soil surface. These communities create an almost living ‘skin’ of intimate associations between soil particles and photoautotrophic (cyanobacteria, algae, lichens, bryophytes) and heterotrophic (bacteria, fungi, archaea) organisms (Weber et al. 2022). Biological soil crusts regularly experience low water activity and are hence expected to harbour halophilic, salt-loving, and halotolerant taxa. These biological crusts prevent soil erosion and retain moisture, properties relevant in desert regions where bioremediation of biological soil crusts could be particularly effective [7].

Permafrost soil remains frozen for at least two consecutive years and is covered by the active layer, i.e., soil that thaws during summer. Despite the frozen conditions, permafrost soils are home to a large diversity of microorganisms [8]. In permafrost and other frozen environments, liquid brine inclusions (cryopegs) form due to salt fractionation [9]. In contrast to the bulk permafrost, nutrients and microorganisms can move in the brine, enabling microbial activity. These brines can reach salt concentrations near saturation, but nevertheless provide a liquid environment for microorganisms to inhabit [10]. Permafrost soil, and especially the cryopegs within, has been a source of many halophilic microorganisms described in the literature [11–13]. Microorganisms isolated from these environments can serve as models in astrobiology [14], and their enzymes and metabolites have biotechnological and industrial applications [15, 16].

Microbial life in Peary Land soils require special adaptations to the low temperatures, low water activity and low nutrient conditions; including modifications to cellular membranes, proteins, genetic material and metabolism [17]. The extremophiles that thrive under these conditions are known as psychrophiles or psychrotolerant (organisms that grow below 10°C) and xerotolerant (organisms that can grow at water activity below 0.8, where 1 is pure water) [18]. Halophiles are a group of xerotolerants that have evolved to grow under high salt conditions with low water activity [19]. Halophilic microorganisms have previously been isolated from non-Polar deserts such as the Atacama Desert [20], but very little is known about halophilic and halotolerant microorganisms in Polar deserts.

Previous studies have shown that the Greenland Ice Sheet is a promising source of biosynthetic gene clusters [21]. With this study, we aim to extend this potential to different Peary Land soils, whose microbiome remains underexplored and may harbour novel bacterial isolates with potential applications in biotechnology. We hypothesize that i) permafrost soil, biological soil crust and surface soil in Peary Land are inhabited by novel culturable halophilic and halotolerant bacteria, ii) the genomes of these bacteria reveal adaptations to life at low water activity and iii) Northern Greenland represents a potential source of secondary bacterial metabolites.

## MATERIALS AND METHODS

### Sampling sites

The Kap København formation (sampled at N82.341460, W21.256961, 160 meters above sea level, m.a.s.l.) (**Fig. 1a**), consists of a 100-m thick succession of shallow marine sediments which were deposited during an interglacial period 2 million years ago. The deposits were later uplifted and exposed subaerially about 1 million years ago, at which time the deposits became permafrost [2]. Samples were extracted using a sterilized shovel from the permafrost ca. 4 meter below the surface accessed from the side of the formation, which was exposed due to recent erosion. Within minutes of extraction, the extracted permafrost material was opened using a sterile chisel and subsamples from the freshly exposed material were immediately transferred to sterile plastic tubes where they were kept until initiation of the isolation work at our laboratories.

**Figure 1.**
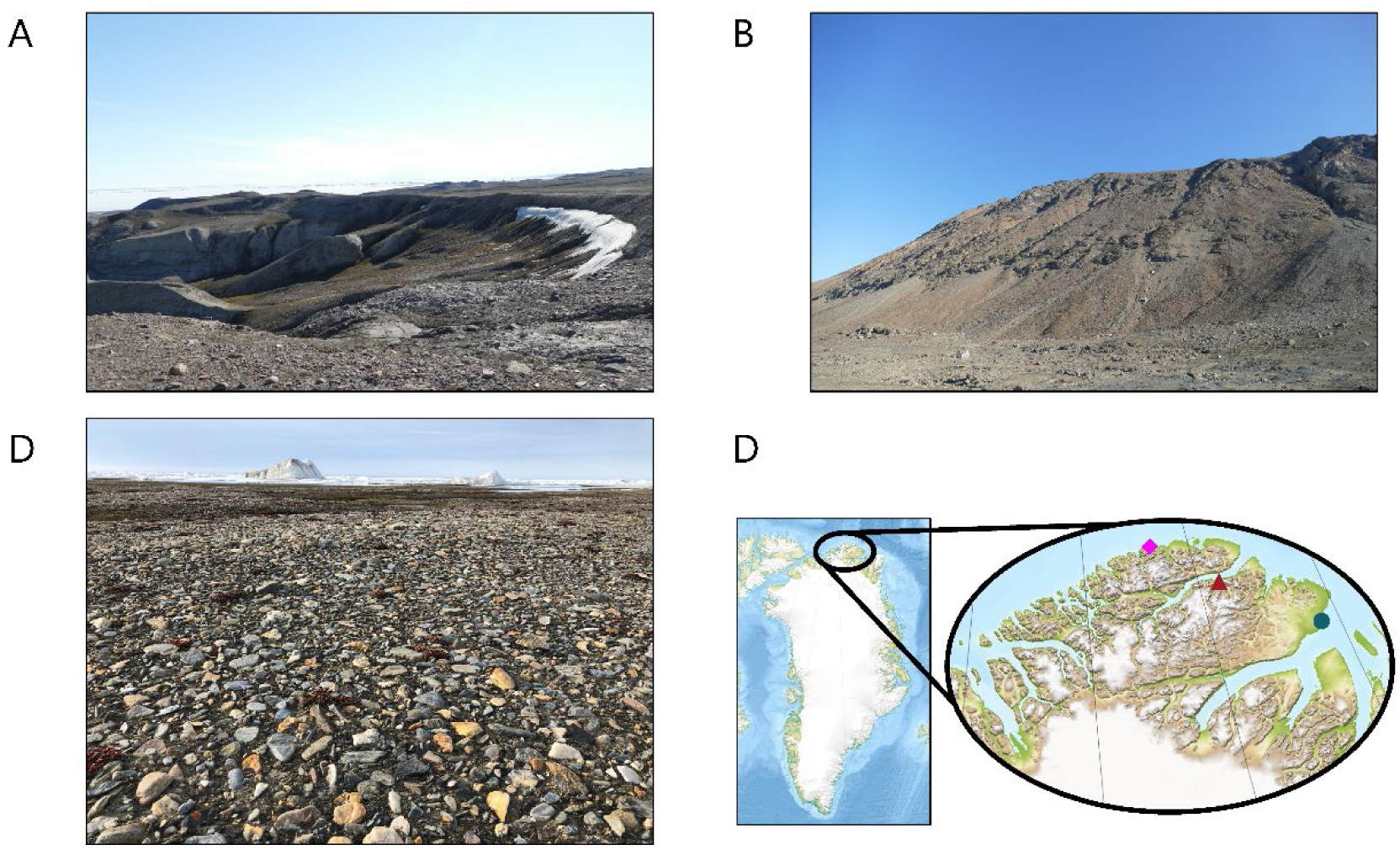
Sampling sites at Kap København (a) where permafrost soil samples were obtained at the erosion front left of the snowpack, Citronen Fjord (b) and Cape Morris Jessup (c). d: Location of Peary Land within Greenland and sampling sites: Cape Morris Jessup (magenta diamond), Citronen Fjord (red triangle) and Kap København (dark green circle). Greenland map: Uwe Dedering, CC BY-SA 3.0. Peary Land map: Blue Green Atlas, CC BY 4.0.

At Citronen Fjord (**Fig. 1b**), conditions are desert-like with very sparse vegetation consisting of scattered specimens of Arctic Willow (*Salix arctica*) and Entire-leaved Mountain Avens (*Dryas integrifolia*). Here, samples of biological soil crust were obtained from a site devoid of vegetation (N83.085782, W28.2496755, 28 m.a.s.l.). Biological soil crust was scooped from the surface using a sterile spoon and transferred to sterile plastic tubes.

Due to its proximity to the Arctic Ocean, Cape Morris Jessup (**Fig. 1c**) harbours more dense vegetation (5-15 percent ground cover) compared to Citronen Fjord. The only weather station in Peary Land, WMO 04301 in Cape Morris Jessup, reported a mean temperature of -14°C, mean annual precipitation of 234 mm, and temperature maxima of around 12°C in the summer for the period 2020-2023 [22]. The vegetation is dominated by Purple Saxifrage (*Saxifraga oppositifolia*) and Arctic Willow. At N83.657236, W33.368495, 5 m.a.s.l., we sampled biological soil crust (as described above) and soil from 0-4 cm depth by hammering sterile steel cores (4.5 cm diameter) into the soil. A subsample from each soil core was transferred to sterile plastic tubes.

### Strain isolation

Samples were stored at 4°C for a period between one and three months, which included the travel time from the sample sites in Peary Land to Copenhagen, Denmark, and pre-incubation of the samples, known as cold enrichment, which has been shown to increase the number of isolated bacteria from permafrost [23].

After pre-incubation, the samples were plated on agar plates. Under sterile conditions, approximately 1 gram of soil was suspended in 1 mL of phosphate-buffered saline (PBS) solution supplemented with 10% NaCl. This buffer was chosen to initially select for bacteria that were resistant to shock from high osmolarity. After gentle mixing for 5 minutes, the samples were briefly centrifuged to remove solid particles from suspension. 100µL of the supernatant were plated on agar plates (see below) and left to grow. For the permafrost samples, the supernatant was diluted 100 and 1000 times prior to plating. The plates were incubated at either 4°C, 10°C or 25°C, and checked weekly and daily, respectively. New colonies were randomly selected based on unique morphologies, then plated into fresh agar plates and incubated again. After incubation, a single colony was picked, plated and incubated. This process was repeated several times until only a single type of colony was observed, at which point we considered that the culture was axenic. Single colonies from the axenic plates were transferred into liquid medium and incubated overnight. 1mL of these cultures was mixed with a glycerol stock (20% v/v final concentration) and frozen at -70°C for long-term storage.

For isolation of halotolerant and halophilic bacteria, three saline media were used. The first was a modified HM medium [24], referred to in this manuscript HM10 for its 10% w/v content of NaCl, and containing, in g L^-1^: NaCl (100) MgSO_4_·7H_2_O (1), CaCl_2_·2H_2_O (0,36), KCl (2), KBr (0,27), FeCl_3_·6H_2_O (0,002), NaHCO_3_ (0,06), Proteose-peptone no. 3 (5), yeast extract (10), glucose (1) and Bacto agar (20, when applicable) adjusted to pH 7,8-8. The second was a medium devised for this study named OHAM (Oligotrophic Halophilic Actinobacteria Medium) containing, in g L^-1^: soluble starch (20), KNO_3_ (0.2), NaCl (100), MgSO_4_·7H_2_O (0.5), K_2_HPO_4_ (0.5), FeSO_4_·7H_2_O (0.01), sodium pyruvate (0.05) and Bacto agar (15, when applicable), and a final pH of 7.2. The third was R2A medium (Sigma Aldritch, Germany) supplemented with 10% NaCl.

In addition, a non-saline medium, Gauze Medium 1 (GM1), was also used, containing in g/L: Soluble starch (20), KNO_3_ (1), NaCl (0.5), MgSO_4_·7H_2_O (0.5), K_2_HPO_4_ (0.5), FeSO_4_·7H_2_O (0.01) and bacto agar (15, when applicable). Final pH was adjusted to 7.2-7.4. This medium has been used in previous works to isolate bacterial strains from permafrost [25].

Premade marine agar (MarA, Sigma Aldritch, Germany) and Tryptic Soy Broth (TSA, Sigma Aldritch, Germany) were used for routine culturing of the isolates after axenic cultures were obtained.

### Determination of the effect of NaCl on growth

To test the NaCl tolerance of the selected isolates, 96-well plates were prepared with HM in different concentrations of NaCl, from 0% (w/v) to 20% in increments of 2.5%. The plates were inoculated with overnight cultures to a constant initial OD_600_, blanks were made for each different NaCl concentration. The plates were then incubated at 25°C in a BioTek Synergy H1 Multimode Reader (Agilent, USA) for 5 days with constant rotation and OD measurements every 30 minutes. Range of growth was determined based on the observed growth in the wells at the end of the experiment. To calculate the effect of NaCl on growth, blanks were subtracted and each replicate was fit to a Gompertz model using the R package *growthrates* [26] version 0.8.4. The resulting parameters were used to calculate the area under the curve using the R package *pracma* [27] version 2.4.4.

### 16S rRNA gene sequencing and analysis

Total DNA was isolated from overnight cultures using the DNeasy PowerSoil Pro Kit (Qiagen, Germany). The 16S rRNA gene was then amplified with the PCRBio kit (PCR Biosystems, UK) using the universal primers 27F (5’-AGA GTT TGA TCM TGG CTC AG-3’) and 1492R (5’-ACG GYT ACC TTG TTA CGA CTT-3’) purchased from TAG Copenhagen (www.tagc.com). After the size and quality of the amplicons was assessed with electrophoresis on agarose gel and ethidium bromide staining, the amplified product was sent for Sanger sequencing to Eurofins (Germany) with the same primers, to obtain both forward and reverse reads.

The forward and reverse reads were aligned in MEGA11 [28] using the MUSCLE algorithm [29] to produce the near full-length 16S rRNA gene sequences. Then, BlastN [30] was used to identify the strains to the genus level; this search was restricted to type strain sequences. To study phylogenetic relationships, an alignment file was produced in MEGA11 using the MUSCLE algorithm with the near full-length 16S rRNA gene sequences and the full-length 16S rRNA gene sequences of related type strains. A phylogenetic tree was constructed with this alignment file using IQ-TREE 2 [31] with ModelFinder [32] and ultrafast bootstrap [33]. Lastly, the tree was drawn with FigTree v1.4.4 (http://tree.bio.ed.ac.uk/software/figtree/) and edited using Inkscape v 1.3.2 (https://inkscape.org/).

### Genome sequencing and assembly

Total genomic DNA was isolated from overnight cultures using the DNeasy PowerSoil Pro Kit (Qiagen, Germany). Short read libraries were prepared using the DNA-Seq Enz kit for MagicPrep (Tecan, USA). The quality of the libraries was checked using a 5200 Fragment Analyzer System (Agilent, USA) before sequencing of the DNA fragments using Miseq2 (Illumina, Germany). The quality of the reads was examined with FastQC v.0.12.0 (https://www.bioinformatics.babraham.ac.uk/projects/fastqc/) and Trimmomatic v0.5.1 [34] was used to remove adapter sequences and low quality reads. Unicycler v0.5.1 [35] was used with SPAdes v4.0.0 [36] to assemble the genomes. The basic statistics and quality of the assemblies were determined with QUAST v5.2 [37], checkM v1.2.3 [38] and BUSCO v5.7.1 [39]. To sequence the genome of *Arthrobacter halodurans* DSM 21081^T^, genomic DNA was directly ordered from the German Collection of Microorganisms and Cell Cultures (DSMZ) and sequenced as described above.

### Genome analysis

The phylogeny of the assembled genomes was determined with the Type (Strain) Genome Server (TYGS) at DSMZ (tygs.dsmz.de), with nomenclature provided by the List of Prokaryotic Names with Standing in Nomenclature (LPSN) [40, 41]. TYGS calculates DNA-DNA-Hybridation (dDDH) distances to the closest type strains of the query genome using the Genome-to-Genome Distance Calculator d_4_ formula (GGDC formula 2). The strains were assigned to the genus of the closest type strain, and they were assigned to a species if dDDH distance was above 70% [42].

The genomes were annotated with Bakta [43] and their metabolic pathways were explored by obtaining KEGG Orthology (KO) numbers using the BlastKOALA and Reconstruct tools of the Kyoto Encyclopedia of Genes and Genomes (KEGG) [44]. AntiSMASH [45] in relaxed mode was used to find biosynthetic gene clusters (BGCs) in the assemblies.

## RESULTS AND DISCUSSION

### Bacterial isolates

A total of 49 bacterial isolates were obtained from the samples (**Table 1**). The soil samples from Citronen Fjord (CF) and Kap København (KK) were the most prolific, while a lower number of isolates was obtained from Cape Morris Jessup soil (CMS) and biological crust (CMC). In addition to bacterial colonies, fungal colonies were observed in plates from all samples. We focused on the bacterial isolates and fungal isolates were not further investigated.

**Table 1.** Number of bacterial isolates obtained from each site and medium. The total number of bacterial isolates was 49. Saline media are marked with an asterisk (*).

| Medium | Citronen Fjord<br>(biological crust) | Cape Morris<br>Jessup (soil) | Cape Morris<br>Jessup (biological<br>crust) | Kap København<br>(permafrost soil) |
| --- | --- | --- | --- | --- |
| GM1 | 10 | 4 | 2 | 16 |
| HM10* | 7 | 0 | 0 | 0 |
| R2A+10% NaCl* | 4 | 2 | 0 | 0 |
| OHAM* | 2 | 0 | 0 | 2 |
| Total | 23 | 6 | 2 | 18 |

The 17 strains isolated with saline media were capable of growth under a wide range of salt concentrations and showed different responses to salt (**Fig. 2**). Three of these isolates, KK5.5, CF3.5 and CF4.19 showed the optimal growth at 0% w/v NaCl, while other isolates such as CF1.1, CF4.1, CF4.2 or CMS1.3 showed almost or no growth without NaCl. The growth of all isolates was impaired at high salt concentrations, but eight isolates showed some level of growth at the maximum concentration tested, 20% w/v NaCl. Interestingly, some isolates belonging to the same genus, e.g., *Nesterenkonia*, showed similar responses to NaCl, while both *Arthrobacter* isolates, KK5.5 and KK5.7, showed slightly different growth patterns as NaCl concentrations increased. Halophiles are commonly classified as such if their growth optimum is above 2% w/v NaCl, while halotolerant organisms grow optimally below 2% and can tolerate a wide range of salinities [46]. Based on this classification, the tested isolates consisted of a mixture of halotolerant (CF3.5, CF4.19 and KK5.5) and halophiles (the rest of the strains).

**Figure 2.**
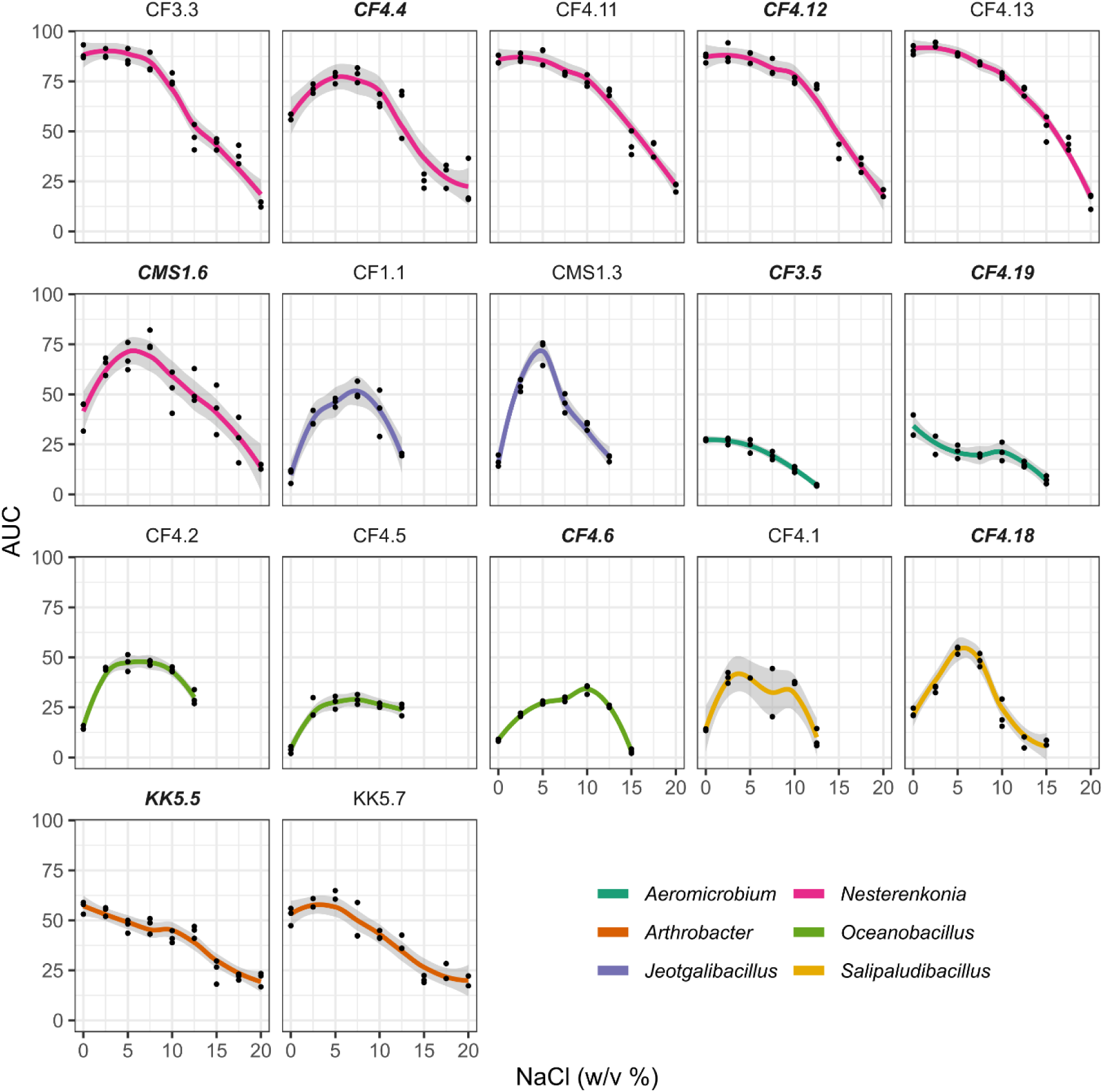
Growth of strains isolated in saline media at 0 to 20% w/v NaCl. Concentrations with no data indicate no measurable growth. Y axis is Area Under the Curve of a Gompertz model fitted to the data to indicate total growth at 72 hours. Black dots represent datapoints. Lines indicate LOESS fit to data with a span of 0.75. Shaded area is the 95% confidence interval. Colour of lines indicates genus. Isolates whose genomes were sequenced are highlighted in bold and italics.

### 16S rRNA gene sequencing

Based on their 16S rRNA gene sequence, the affiliation of the isolates with known type strains was visualized (**Fig. 3**). Full details of each isolate and accession numbers are available in Table S1. The isolates mainly belonged to three phyla: Actinomycetota, Bacillota and Pseudomonadota, while Bacteroidota was only represented by one isolate, *Acidovorax* sp. CMS1.1 (Fig. 3B). All isolates demonstrated the ability to grow at 4°C. Many of the strains were isolated in saline media, thus demonstrating their halotolerance. These included all the Bacillota, as well as isolates belonging to the genus *Nesterenkonia*. Some strains isolated in non-saline media also proved to be halotolerant, as they grew on MarA (containing 1.945% w/v NaCl), including most members of *Arthrobacter* and *Pseudomonas* sp. KK6.1. However, other isolates, like *Massilia* sp. CMS3.1 and *Flavobacterium* sp. CMS1.2 were unable to grow in medium supplemented with >1% NaCl (data not shown).

**Figure 3.**
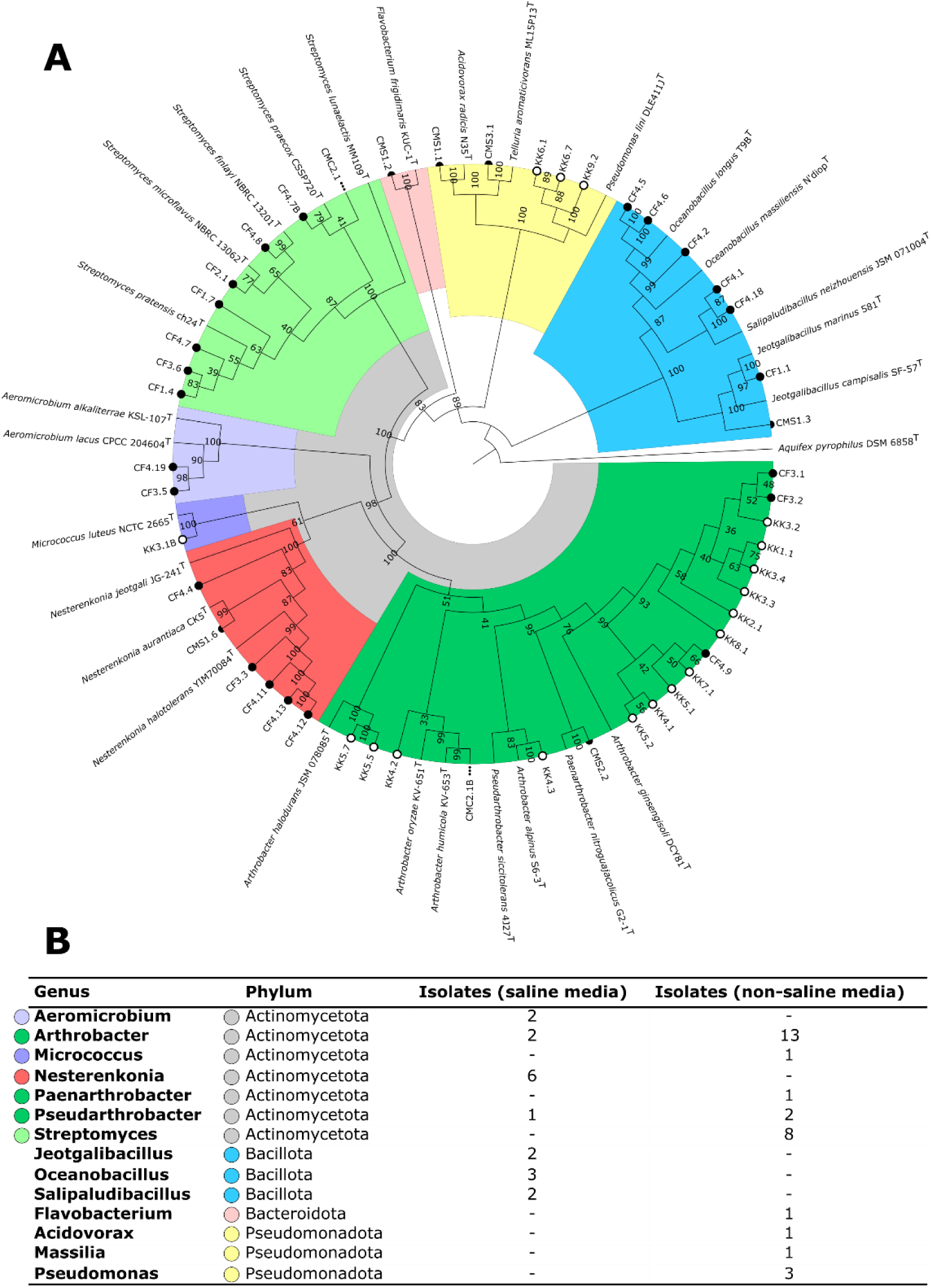
Phylogenetic affiliation of isolates. **A**: Consensus cladogram based on the 16S rRNA gene sequences of the isolates and selected type strains calculated with TIM2+F+I+G4. Bootstrap values (as percentage of 10000 repetitions) are shown at the branch points. Phyla are highlighted with colours as indicated in panel B. Within Actinomycetota, genera are further highlighted with colours as indicated in panel B. The markers indicate the source of isolation: Citronen Fjord (black circle), Kap København (white circle), and Cape Morris Jessup soil (semicircle), and crust (dots). Type strains have no markers. Aquifex pyrophilus DSM 6858 was used as the outgroup. **B**: Number of isolates of each detected genus, and colours used to highlight groups in panel A.

The most abundant genera were *Arthrobacter*, predominantly isolated from permafrost soil, and *Streptomyces* and *Nesterenkonia*, obtained from biological crust. Previous studies have shown that these genera, as well as other genera isolated in this study, are commonly found in polar environments. For example, Singh and colleagues [47] reported that *Arthrobacter* and *Pseudomonas* were the most commonly isolated genera in permafrost soil from Svalbard, Norway, where they also isolated strains belonging to *Nesterenkonia, Bacillus*, and others. Interestingly, the authors also obtained isolates belonging to *Streptomyces*, while none of our isolates in this genus were obtained from permafrost soil. *Streptomyces* and other Actinomycetota are commonly found in arid and desert soils [48] as well as polar environments [49], and extremophilic and extremotolerant *Streptomyces* have drawn attention as potential producers of novel antibacterial compounds and other compounds of interest [50]. In addition, *Nesterenkonia* has been isolated from Arctic permafrost soil [51], as well as other cold environments, like soils of Antarctica and the Tibetan plateau [52, 53].

*Arthrobacter* is commonly found in permafrost soil with both culture-dependent [47, 54] and culture-independent methods [55], the latter of which have shown that *Arthrobacter spp.* can constitute up to 4% of the total bacterial community [56]. This suggests that either *Arthrobacter spp.* are especially adapted to survive in permafrost soil, or that they can quickly adapt to newly forming permafrost. The *Arthrobacter* pangenome [57] revealed great potential adaptability, which could explain its prevalence in permafrost soil. The same study showed the halotolerance of several *Arthrobacter* strains, a similar observation to our *Arthrobacter* isolates, as many of them could grow on MarA or were directly isolated in saline medium. Based on its prevalence in a variety of environments, some authors have used *Arthrobacter* as a model to study bacterial adaptations in soil [55, 58]. In our study, the isolated *Arthrobacter* strains were initially isolated in either GM1 or OHAM, media with no nitrogen source other than nitrate, and with glycogen as the only carbon source (along with pyruvate in OHAM). Some members of this genus are known to be able to assimilate nitrate and nitrite for growth [59, 60]. Based on our results, GM1 may be an effective selective medium to isolate *Arthrobacter* strains from permafrost soil.

The diversity of isolates obtained from Kap København, Cape Morris Jessup and Citronen Fjord, and the displayed halotolerance of the isolates, show that the sampled sites are inhabited by a wide range of halotolerant bacteria, thus partially failing to reject our first hypothesis. In the next section, we discuss the novelty of the isolates.

### Genomes and functional analyses of selected isolates

From the 16S rRNA gene-based phylogeny, nine isolates were selected for whole genome analysis (**Table 2**), of which eight were halotolerant. *Massilia* sp. CMS3.1 was determined to not be halotolerant, as it was only able to grow in GM1 with <1% NaCl, but was chosen for genome sequencing (a) as a Gram-negative and non-halotolerant representative and (b) to investigate its adaptations to its natural environment as a non-halotolerant bacterium. In the case of *Arthrobacter* sp. KK5.5, its closest strain was *A. halodurans* DSM 21081^T^, whose genome was not available in any database. To confirm the phylogeny of our isolate, we sequenced the genome of *A. halodurans* DSM 21081^T^. All genomes showed completeness above 98% and contamination below 2%, which was determined sufficient to proceed to the next part of the analysis. Six of the nine sequenced isolates showed dDDH similarities to their closest relatives below 30%, well below the established threshold of 70% to distinguish between different species (**Table 2**). Following the phylogenetic analysis, we conducted a functional analysis of the genomes using the KEGG database and KEGG Orthology numbers

**Table 2.**
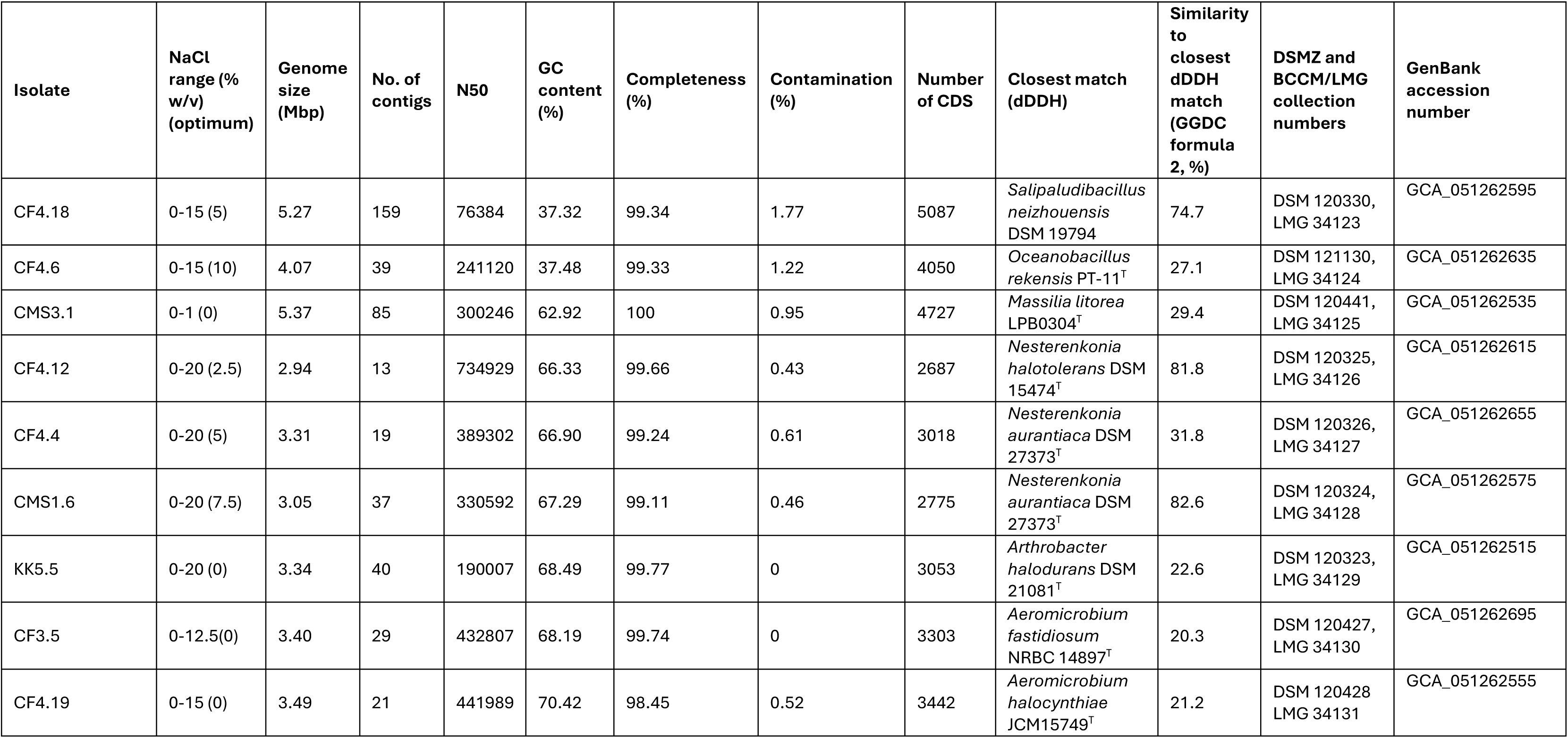
Isolates selected for genome sequencing, information on the assemblies, and similarity to their closest relatives bases on digital DNA-DNA hybridization (dDDH). Completeness and contamination were calculated with CheckM. Completeness calculated by BUSCO is available in Table S2. GGDC: Genome-to-Genome Distance Calculator. DSMZ: German Collection of Microorganisms and Cell Cultures. BCCM/LMG: Belgian Co-ordinated Collections of Micro-organisms / Laboratory of Microbiology, Ghent University.

#### Adaptations to high salinity and low water activity

All sequenced isolates, except for *Massilia* sp. CMS3.1, possessed orthologs of genes encoding importers of osmoprotectants like proline, ectoine and betaine choline (**Fig. 4** and **Table S3**). Osmoprotectants are small, neutral organic solutes that do not interfere with cellular processes, and can thus be accumulated to mitigate osmotic and low-temperature stress [61]. The ability to import osmoprotectants from the environment offers flexibility, allowing bacteria to adapt to fluctuating salinity. This strategy reduces the energy cost associated with de novo synthesis, providing a competitive advantage. Interestingly, the non-halotolerant *Massilia* sp. CMS3.1 possessed only one ortholog of osmoprotectant transporters (*yaaJ*), whereas its close relative, *Telluria aromaticivorans*, has 16 transporters and 9 synthesis-related genes, despite its low salt tolerance [62]. Conversely, *Salipaludibacillus neizhouensis* CF4.18 had 33 orthologues related to osmoprotectant transport, almost twice that of the type strain of its species, revealing large intra-species variation. These findings suggests that halotolerance may not be directly corelated with the copy number of genes related to osmoprotectant synthesis and import.

**Figure 4.**
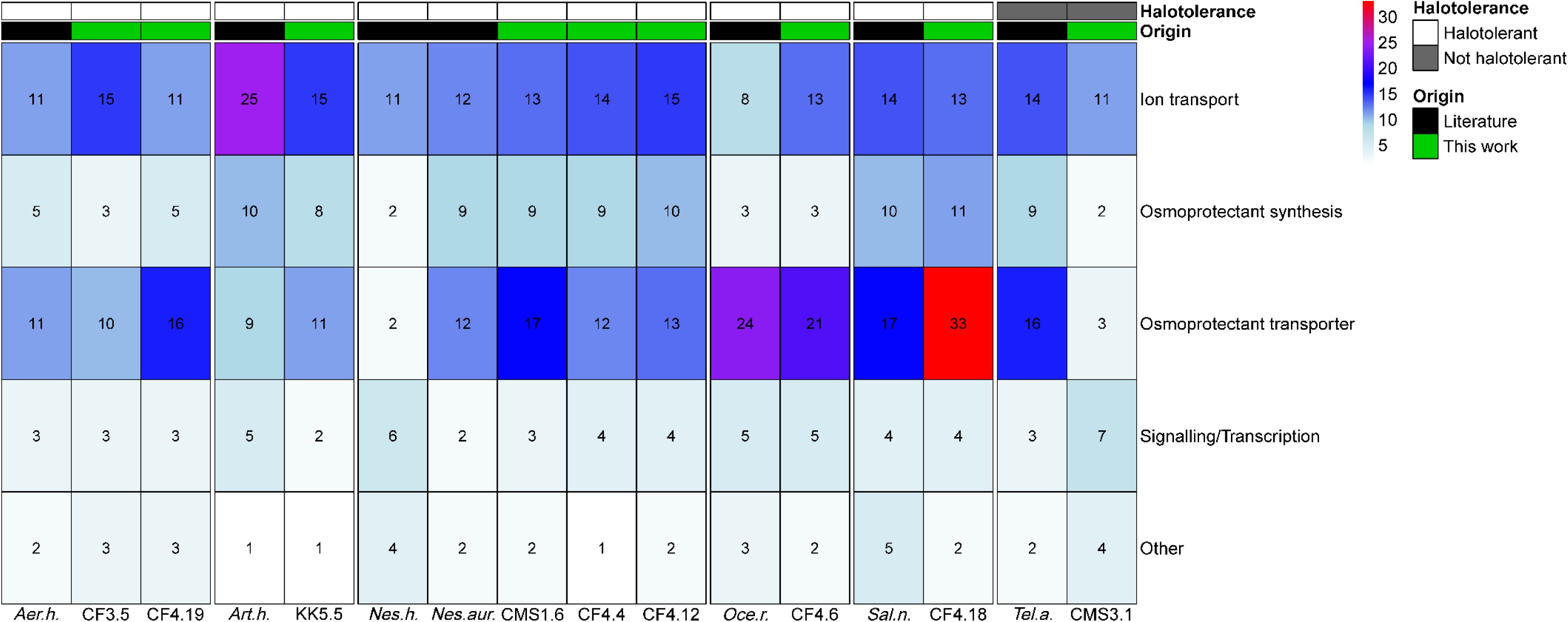
KEGG Orthology count numbers of groups of genes that have been described to be important for halotolerance in the sequenced genomes and in genomes of close relatives. The complete dataset is available in Table S3. The close relatives are as follows, from left to right: *Aer. h., Aeromicrobium halocynthiae* JCM 15749^T^; *Art. h., Arthrobacter halodurans* DSM 21081^T^; *Nes. h., Nesterenkonia halotolerans* DSM 15474^T^; *Nes. aur., Nesterenkonia aurantiaca* DSM 27373^T^; *Oce. r., Oceanobacillus rekensis* PT11^T^; *Sal. n., Salipaludibacillus neizhouensis* DSM 19794^T^; *Tel. a., Telluria aromaticivorans* ML15P13^T^.

Other abundant groups were orthologues related to ion transport (including *trkA*, *kdpABC*, *tlyC*, and *nhaAC*). Ion transporters are very common in bacteria and are not necessarily indicative of halotolerance, as evidenced by similar numbers of ion transporter orthologues in non-halotolerant strains (**Fig. 4**). However, ion transporters may have specific adaptations that aid with halotolerance. For example, the cyanobacterium *Aphanothece halophytica* has ion transporters with different ion specificities that allow it to tolerate high salinity and pH [63]. In the halophilic bacterium *Halomonas beimenensis*, the genes *nqrA*, *trkA*, *atpC*, *nadA*, and *gdhB* — related to ion transport — along with *spoT*, *prkA*, *mtnN*, *rsbV*, *lon*, *smpB*, *rfbC*, *rfbP*, *tatB*, *acrR1*, and *lacA —* related to cellular signalling, quorum sensing, transcription/translation, and cell motility — are critical for halotolerance, and their deletion impairs growth at high salinity [64]. Similarly, the genes *trkA2*, *smpB*, *nadA*, *mtnN2*, *rfbP*, *lon*, and *atpC* are upregulated in high salinity in *Virgibacillus chiguensis* [65]. Orthologues of many of these genes were found in varying numbers in the sequenced isolates (**Table S3**), suggesting their potential role in the observed halotolerance across the bacteria isolated in this study.

Cold temperatures can lead to similar stress variables than high salinity. For example, water activity is reduced as ice crystals form, leading to osmotic stress, and brines can form as salt ions are excluded from ice crystals [9]. Thus, the halotolerance-related genes discussed may also aid the isolates against the low temperatures in their native Northern Greenland. In addition, the genomes also contained several orthologues of cold shock proteins (*cspA*). These proteins have a protective effect against the stresses associated with low temperatures by binding to single stranded nucleic acids and regulating transcription and translation [66].

#### Other metabolic pathways

Some of the sequenced genomes contained genes encoding enzymes related to nitrogen and sulphur cycling. *S. neizhouensis CF*4.18 and *Oceanobacillus* sp. CF4.6 had a complete assimilatory sulfate reduction pathway, while *N. aurantiaca* CMS1.6 and *N. halotolerans* CF4.12 lacked this pathway completely. The rest of the sequenced isolates had an incomplete pathway, lacking the enzymes to reduce sulphate to sulphite. However, all genomes contained transporters for alkanesulfonates, and most genomes contained genes encoding for alkanesulfonate monooxigenases. This alternative source of sulphur may aid against sulphur starvation [67], as the low amount of organic carbon in Peary Land soils could limit the amount of sulphur available from organic matter [3]. However, sulphur may not be a limiting nutrient in the permafrost at Kap København, as it consists mostly of marine deposits, which are typically enriched in sulphur [68, 69].

*Nesterenkonia* sp. CF4.4, *Aeromicrobium* sp. CF3.5, *Arthrobacter* sp. KK5.5, *S. neizhouensis* CF4.18 and *Massilia* sp. CMS3.1 had genes encoding for both nitrate and nitrate reductases, while these were not found in the other isolates. Interestingly, *Oceanobacillus* sp. CF4.6 was the only isolate whose genome contained *nirK*, which encodes for a nitric oxide-producing nitrite reductase. The reduction of nitrite to nitric oxide (NO) is the second step in denitrification after the reduction of nitrate to nitrite. NO is then reduced to nitrous oxide (N_2_O), which is finally reduced to N_2_. NO and N_2_O have significant impact on atmospheric chemistry, as they can induce the formation of tropospheric ozone and form NO_X_; furthermore, N_2_O is a potent greenhouse gas. In arid areas, especially in the polar regions, nitrification and denitrification remain poorly understood [70]. There are no available studies on the nitrification or denitrification rates, or on the emission of NO and N_2_O, from Northern Greenland, but ecosystem N losses from Arctic soils through N_2_O emission are generally small, which may be linked to the N limitation of these soils [71, 72]. *Oceanobacillus* sp. CF4.6 may play a role in denitrification and ecosystem loss of nitrogen in its native environment, although environmental studies are needed to characterise the soil nitrogen cycle in the High Arctic desert of Northern Greenland.

All the sequenced genomes encoded pathways for the synthesis of terpenoid backbones. Apart from *Oceanobacillus* sp. CF4.6, whose genome encoded for the mevalonate pathway, the sequenced genomes encoded for the MEP/DOXP pathway for isoprenoid backbone synthesis. Interestingly, the pathway in *N. halotolerans* CF4.12 is incomplete, as it is missing 2-C-methyl-D-erythritol 4-phosphate cytidylyltransferase (EC 2.7.7.60) encoded by *IspD*, and 2-C-methyl-D-erythritol 2,4-cyclodiphosphate synthase (EC 4.6.1.12), encoded by *IspF* [73]. *N. halotolerants* DSM 15474^T^ shows the same missing enzymes in its draft genome. These findings suggest that *N. halotolerans* may contain undescribed isoforms of these enzymes. The synthesis of terpenoid backbones is an important pathway to produce essential and secondary metabolites like undecaprenol phosphate and carotenoids. In fact, all isolates apart from *Oceanobacillus* sp. CF4.6 and the two *Aeromicrobium* sp. had BGCs encoding for terpene-related products, with many of them encoding for carotenoids (see next section). This pathway may be of particular relevance in *Nesterenkonia*, as all isolates belonging to this genus had a bright red or orange colour, suggesting a significant content of carotenoids.

The adaptations discussed in this section allow the isolates to sustain activity in their cold, nutrient-poor, and water-scarce environment. Furthermore, most of the sequenced genomes showed very low dDDH percentages towards their closest known relative, suggesting that they represent previously undescribed species. Thus, we fail to reject hypotheses i and ii.

### Biosynthetic gene clusters in the genomes of the sequenced isolates

The sequenced genomes were found to contain several BGCs (**Fig. 5** and **Tables S4-S12**), including several with no hits to known BGCs. 32% of the identified BGCs putatively encoded for antimicrobial compounds, such as the lassopeptide paeninodin found in *S. neizhouensis* CF4.18 and *Oceanobacillus* sp. CF4.6, ε-Poly-L-lysine synthesis clusters found in *N. halotolerans* CF4.12 and *Nesterenkonia* sp. CF4.4, and other BGCs encoding for lactones found in several isolates. Most of these putatively antimicrobial clusters encoded for polyketide synthases (PKS) and non-ribosomal peptide synthases (NRPS). Antimicrobial compounds and antimicrobial resistance are widely distributed in soils, where competition for available space and resources exerts significant selective pressure [74]. The production of antimicrobial compounds typically incurs a high metabolic cost; for example, Schlatter and Kinkel [75] found trade-offs between efficient growth, antimicrobial resistance and antimicrobial production in *Streptomyces spp*., and suggested that a “super-killer” phenotype results in a reduction in niche width. The most common strategy in *Streptomyces spp.* was the simultaneous production of different kinds of antimicrobial compounds to increase their synergistic effect and slow down the evolution of resistance. However, in carbon-poor soils, as is the case in Peary Land, nutrient limitation may present an obstacle for this strategy. On the other hand, nutrient scarcity may increase evolutionary pressure to inhibit potential competitors. Many studies have demonstrated the production of antibacterial and antifungal compounds by bacteria isolated from arid soils (e.g., Saadoun et al. 2008; Nasfi et al. 2018), and even the hyper-arid Atacama desert is a potential source of novel antimicrobials [48]. Polar desert soils, although not as explored, have also demonstrated this potential [78]. In permafrost, where biomass can be higher and environmental conditions are stable, antibiotic resistance is widespread, even in ancient permafrost [79, 80]. Although measuring the production of antimicrobial compounds of our isolates was beyond the scope of this study, the diversity of BGCs encoding for antimicrobial compounds found in the sequenced genomes suggest a competitive environment where it may be advantageous to have different antimicrobial strategies available.

**Figure 5.**
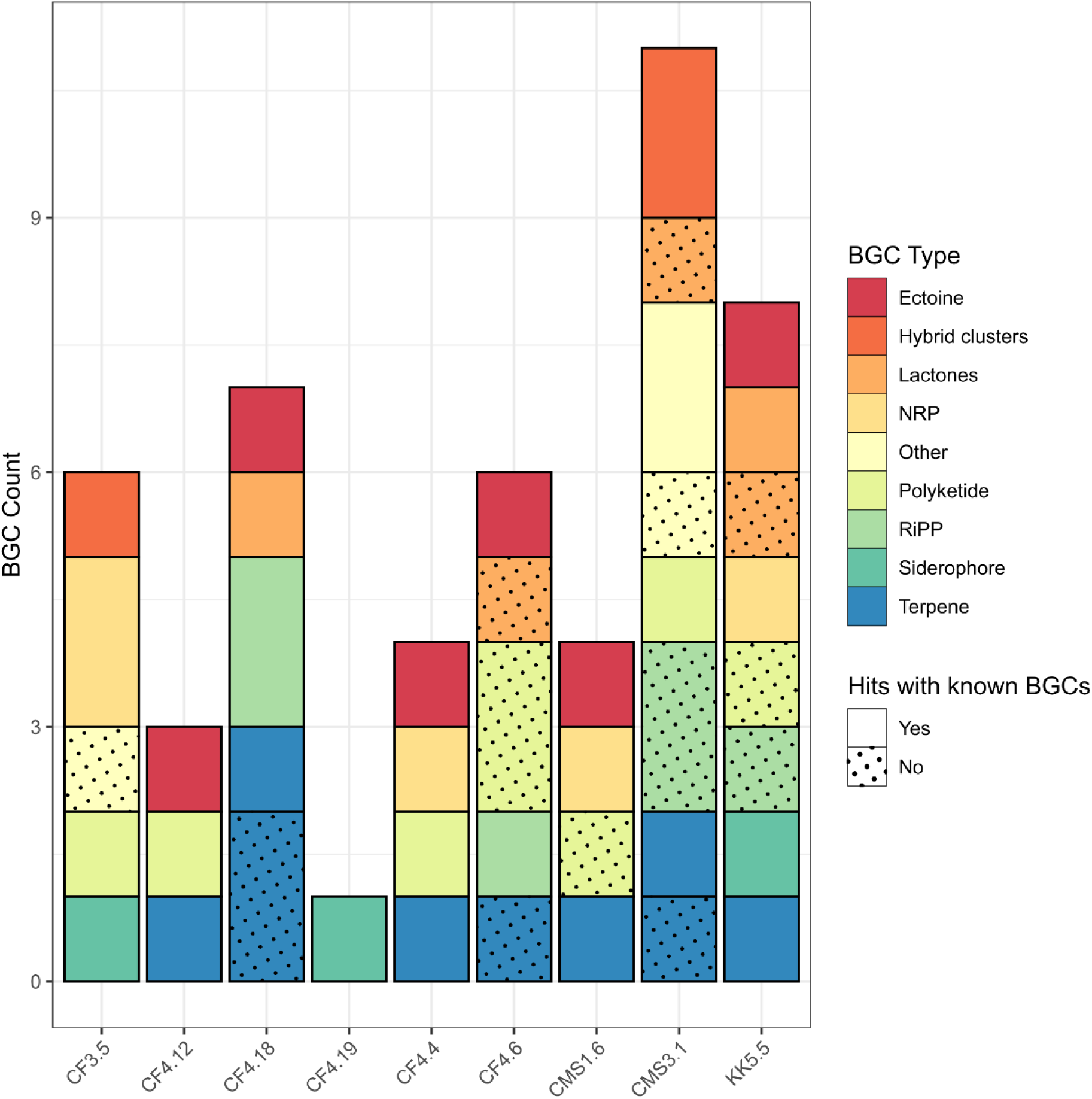
Biosynthetic Gene Clusters (BGCs) found in the assembled genomes. The dotted pattern represents BGCs with no hits to known BGCs. NRP: Non-Ribosomal Peptide. RiPP: Ribosomally synthesized and Post-translationally modified Peptide.

Desert soils often have higher pH compared to soils from forests and other biomes. The high pH is caused by accumulation of minerals and salts that are not washed away by rain. Peary Land soil is no exception, with soils at Cape Morris Jesup and Citronen Fjord having pH of 7.8 and 9.2, respectively. At these pH values, the bioavailability of many metals is lowered. Thus, bacteria in desert soils need effective metal-acquiring systems such as siderophores. Siderophores allow for the solubilisation of iron essential for the production of, e.g., heme proteins. Both *Aeromicrobium* isolates, as well as *Arthrobacter* sp. KK5.5 had BGCs encoding for NRPS-independent siderophores; their genomes also contained orthologs of *fepBDG*, which encode for Fe-siderophore complex transporters, and both *Aeromicrobium* isolates additionally had *fepC*. Notably, *N. halotolerans* CF4.12 also had a complete set of *fepBCDG*, while the other two *Nesterenkonia* isolates had orthologues of *fepCG*. This absence of siderophore synthesis genes, while transport genes are present, may be due to a variety of causes. One possibility is that these bacteria rely on siderophores produced by other bacteria for iron uptake [81].

Lastly, *Massilia* sp. CMS3.1 had a BGC whose putative product is a homoserine lactone. While it had no hits to known clusters, it contained, among others, a protein with an acyltransferase domain; this suggests that this BGC may encode for an N-Acyl homoserine lactone, a group of signalling molecules involved in quorum sensing and biofilm formation. This aligns with the possible ability of *Massilia* sp. CMS3.1 to form biofilms. Indeed, the genome of this isolate contained several genes involved in quorum sensing and biofilm formation, such as orthologues of *lasIR* and *wspBEFR*, as well as *pilGHIJ*, which are involved in type IV pilus formation. Young (<24h) colonies of *Massilia* sp. CMS3.1 were orange, soft and easily pickable, but older (especially >48h) colonies became darker, hard, and difficult to disaggregate, which suggests the formation of a hard extracellular matrix that may be mediated by these genes.

Biofilm formation has been shown to be a successful strategy to improve survivability against extreme environmental factors like temperature and pH (high and low), aridity and high salinity [82]. For example, biofilms are employed by epiphytic bacteria to survive periods of extreme cold and freeze-thaw cycles [83].

Other types of BGCs found in the genomes included an ectoine synthesis cluster and terpenoid synthesis clusters, which were discussed in the previous section; the terpenoid clusters encoded mainly for carotenoids.

It is possible that some BGCs were missed by antiSMASH. Indeed, BGC detection on contig-level assemblies can be challenging, as BGCs may be split into different contigs [84]. This is evidenced by a few BGCs that were found at the beginning or the end of a contig, suggesting that these BGCs are incomplete. This, combined with the significant amount of BGCs that had no hits with known BGCs, suggest that Peary Land is indeed a source of potentially novel secondary metabolites, as per hypothesis iii; however, further investigation *in vivo* will be necessary to fully characterise these secondary metabolites.

## CONCLUSIONS

We isolated 49 bacteria from three different sites in Peary Land, northern Greenland, representing some of the most northernly bacterial isolates, and the first strains isolated from these areas. We showed the diversity of culturable halotolerant bacteria at the three sites and found potentially novel BGCs that suggest a competitive environment with many antimicrobial compounds present. The isolates showed adaptations to their cold and saline native environments, including a variety of ion transporters and genes related to osmoprotectant synthesis and transport. These findings underscore the importance of osmoprotectants in cold and saline environments and expand our understanding of microbial life in extreme environments. Overall, this work not only provides the first insights into the diversity of culturable halotolerant bacteria of northern-most Greenland, but also suggests that the region harbours untapped microbial resources that may hold significant biotechnological value.

## DATA AVAILABILITY

All sequences obtained and used in this study, including near-complete 16S rRNA gene sequences, genome assemblies, and short-read libraries, are available in the National Center for Biotechnology Information (NCBI) database under the BioProject accession number PRJNA1156031.

## ACKNOWLEDGEMENTS

We would like to thank Lorrie Maccario for her help in the library preparation and sequencing of the genomes, and Anette Hørdum Løth and Ayoe Lüchau for their indispensable assistance in the lab. We would also like to thank Magnus Per Damsø Jeppesen for his help in the collection of some of the salt response data. We are grateful to the Greenlandic authorities for granting permission to sample in the Northeast Greenland National Park (Expedition Permit C-21-549) and to export the soil samples to Denmark for scientific work (Non-exclusive license number G21-031 for utilization of Greenlandic genetic resources). We gratefully acknowledge the Ministry of Business, Mineral Resources, Energy, Justice and Gender Equality of the Government of Greenland for granting permission to deposit the strains in DSMZ and BCCM/LMG under conditions that ensure compliance with the International Code of Nomenclature of Prokaryotes.

## FUNDING

This project was funded by the Novo Nordisk Foundation under the NNF Interdisciplinary Synergy Program (grant number NNF19OC0057374): Effects of bacteria on atmospheres of Earth, Mars, and exoplanets -- adapting and identifying life in extraterrestrial environments. This project was also supported by The Danish National Research Foundation within the Center for Volatile Interactions (VOLT, DNRF168). The bacteria in this study were isolated from samples collected during the Leister Around North Greenland 2021 Expedition financed by the Leister Stiftung, Switzerland.

## CONFLICTS OF INTEREST

The authors declare no conflicts of interest.

---

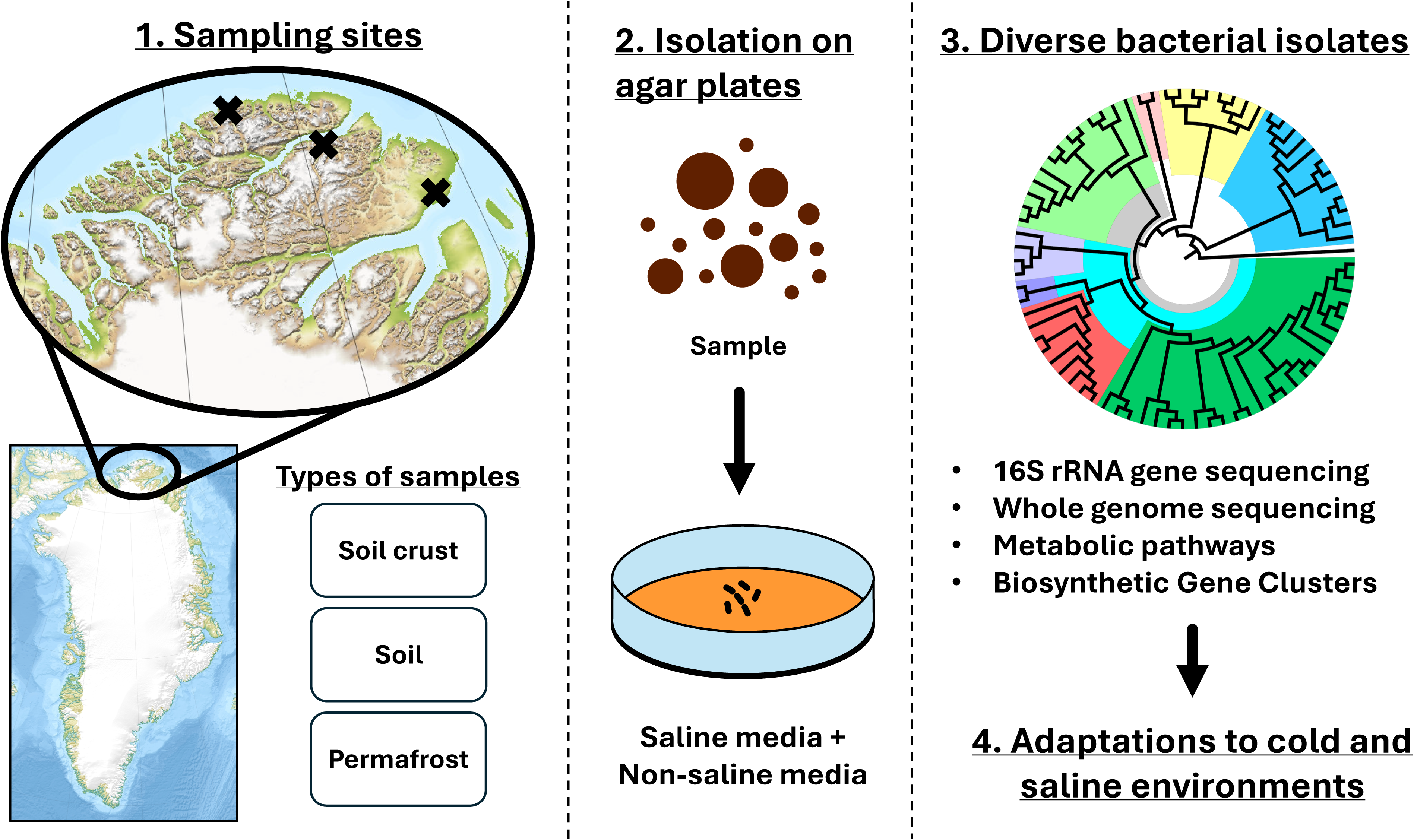

## Notes

### Competing Interest Statement

The authors have declared no competing interest.

### Summary of Updates

Revised NaCl response data for a more thorough analysis. Revised list of genes for Fig. 4. Re-rendered some figures if data was revised or to improve readability. Added strain collection numbers to Table 2.

