## Supplementary data for "Genomic adaptations of novel halotolerant bacteria from extreme North Greenland"

for

Note: The supplementary material also includes a separate .xlsx file with Tables S1 and S3

**Table S1 [separate .xlsx file].** Details on the isolation conditions of each isolate and closest matches to their 16S rRNA gene sequences. The sample of origin is coded into the name of the strain: CF (Citronen Fjord), CMC/CMS (Cap Morris Jessup crust/soil, respectively) and KK (Kap København Formation). Taxonomic classification was based on the closest match to type strains to the genus level. N/S: Not specified. *: The 16S rRNA gene sequence of strain CMS3.1 was closest to *Telluria aromaticivorans* ML15P13$^T$. However, whole genome dDDH indicated that it was closest to *Massilia litorea* LPB0304$^T$, so it was assigned to the genus *Massilia*.

**Table S2.** Genome assembly quality statistics determined by BUSCO (Manni *et al.* 2021).

| **Strain** | **Complete (C)** | **Single-Copy (S)** | **Duplicated (D)** | **Fragmented (F)** | **Missing (M)** | **Total BUSCOs (n)** |
|---|---|---|---|---|---|---|
| *Salipaludibacillus neizhouensis* CF4.18 | 99.5 | 98.4 | 1.1 | 0.0 | 0.5 | 450 |
| *Oceanobacillus* sp. CF4.6 | 100.0 | 99.3 | 0.7 | 0.0 | 0.0 | 450 |
| *Massilia* sp. CMS3.1 | 99.1 | 98.5 | 0.6 | 0.3 | 0.6 | 688 |
| *Nesterenkonia halotolerans* CF4.12 | 98.9 | 98.9 | 0.0 | 0.4 | 0.7 | 537 |
| *Nesterenkonia* sp. CF4.4 | 98.5 | 98.3 | 0.2 | 0.2 | 1.3 | 537 |
| *Nesterenkonia aurantiaca* CMS1.6 | 98.5 | 98.3 | 0.2 | 0.4 | 1.1 | 537 |
| *Arthrobacter* sp. KK5.5 | 100.0 | 100.0 | 0.0 | 0.0 | 0.0 | 537 |
| *Aeromicrobium* sp. CF3.5 | 99.1 | 98.8 | 0.3 | 0.3 | 0.6 | 649 |
| *Aeromicrobium* sp. CF4.19 | 98.6 | 98.6 | 0.0 | 0.3 | 1.1 | 649 |

**Table S3 [Separate .xlsx file].** KEGG orthologues of genes related to halotolerance found in one or more of the isolates. The accession numbers are: *Aeromicrobium halocynthiae* JCM 15749$^T$ (GCA_039531385); *Aeromicrobium* sp. CF 3.5 (GCA_051262695); *Aeromicrobium* sp. CF4.19 (GCA_051262555); *Arthrobacter halodurans* DSM 21081$^T$ (GCA_041877255); *Arthrobacter* sp. KK5.5 (GCA_051262515); *Nesterenkonia halotolerans* DSM 15474$^T$ (GCA_014874065); *Nesterenkonia aurantiaca* DSM 27373$^T$ (GCA_004364585); *Nesterenkonia aurantiaca* CMS1.6 (GCA_051262575); *Nesterenkonia* sp. CF4.4 (GCA_051262655); *Nesterenkonia halotolerans* CF4.12 (GCA_051262615); *Oceanobacillus rekensis* PT 11$^T$ (GCA_002153375); *Oceanobacillus* sp. CF4.6 (GCA_051262635); *Salipaludibacillus neizhouensis* DSM 19794$^T$ (GCA_002886185); *Salipaludibacillus neizhouensis* CF4.18 (GCA_051262595); *Telluria aromaticivorans* ML15P13$^T$ (GCA_013003915); *Massilia* sp. CMS3.1 (GCA_051262535). The genes were selected based on previous studies (Barberán *et al.* 2017; Bowman 2017; Chen *et al.* 2017; Lee and Kim 2022) and the KEGG database. The strains isolated in this study are highlighted in bold and with a darker shade.

**Table S4.** Biosynthetic gene clusters in *Salipaludibacillus neizhouensis* CF4.18.

| Cluster ID | BGC type | Location (bp) | Most similar known cluster | Class | Similarity | Putative function |
|---|---|---|---|---|---|---|
| 4.1 | T3PKS | 126961-168049 | Thioholgamide A | RiPP | 4% | Antimicrobial |
| 6.1 | Terpene | 17227-31739 | Carotenoid | Terpene | 83% | Antioxidant |
| 8.1 | Lassopeptide | 69883-93806 | Paeninodin | RiPP | 80% | Antimicrobial |
| 13.1 | Ectoine | 38061-48447 | Ectoine | Other | 66% | Osmoprotectant |
| 33.1 | Terpene | 22034-42882 | No hits | N/A | N/A | Unknown |
| 53.1 | Terpene | 1-20913 | No hits | N/A | N/A | Unknown |

**Table S5.** Biosynthetic gene clusters in *Oceanobacillus* sp. CF4.6.

| Cluster ID | BGC type | Location (bp) | Most similar known cluster | Class | Similarity | Putative function |
|---|---|---|---|---|---|---|
| 1.1 | T3PKS | 9736-50827 | No hits | N/A | N/A | Unknown |
| 1.2 | T3PKS | 487840-529018 | No hits | N/A | N/A | Unknown |
| 5.1 | Ectoine | 144893-155279 | Ectoine | Other | 66% | Osmoprotectant |
| 7.1 | Lassopeptide | 8181-34394 | Paeninodin | RiPP | 100% | Antimicrobial |
| 9.1 | Terpene | 140128-160946 | No hits | N/A | N/A | Unknown |
| 11.1 | Betalactone | 124195-143348 | No hits | N/A | N/A | Unknown |

**Table S6.** Biosynthetic gene clusters in *Massilia* sp. CMS3.1.

| Cluster ID | BGC type | Location (bp) | Most similar known cluster | Class | Similarity | Putative function |
|---|---|---|---|---|---|---|
| 1.1 | Terpene | 282284-304068 | No hits | N/A | N/A | Unknown |
| 1.2 | Redox cofactor | 320972-343093 | Lankadicin C | NRP+polyketide | 13% | Antimicrobial |
| 1.3 | Acyl amino acids | 362612-423361 | Jerangolid A/D | Polyketide | 9% | Antifungal |
| 1.4 | RiPP-like | 630212-642416 | No hits | N/A | N/A | Unknown |
| 1.5 | Homoserine lactone | 803268-815363 | No hits | N/A | N/A | Signaling molecule |
| 3.1 | NRPS-like | 212059-255652 | luminmycin A/glidobactin A/cepafungin | NRP+Polyketide | 7% | Antimicrobial |
| 5.1 | Terpene | 323256-346934 | Carotenoid | Terpene | 100% | Antioxidant |
| 6.1 | Acyl amino acids | 100196-160981 | N-tetradecanoyl tyrosine | Other | 6% | Antimicrobial |
| 6.2 | NRPS-like | 164525-207245 | K53 capsular polysaccharide | Saccharide | 10% | Cellular defense |
| 9.1 | Indole | 132220-153491 | No hits | N/A | N/A | Signaling molecule |
| 10.1 | RiPP-like | 104067-115713 | No hits | N/A | N/A | Unknown |

**Table S7.** Biosynthetic gene clusters in *Nesterenkonia halotolerans* CF4.12.

| Cluster ID | BGC type | Location (bp) | Most similar known cluster | Class | Similarity | Putative function |
|---|---|---|---|---|---|---|
| 1.1 | Ectoine | 35127-45516 | Ectoine | Other | 50% | Osmoprotectant |
| 1.2 | Terpene | 297796-318671 | Carotenoid | Terpene | 28% | Antioxidant |
| 4.1 | T3KS | 251487-292569 | Oryzanaphthopyran | Polyketide | 10% | Antimicrobial |

**Table S8.** Biosynthetic gene clusters in *Nesterenkonia* sp. CF4.4.

| Cluster ID | BGC type | Location (bp) | Most similar known cluster | Class | Similarity | Putative function |
|---|---|---|---|---|---|---|
| 2.1 | Ectoine | 116122-126520 | Ectoine | Other | 50% | Osmoprotectant |
| 2.2 | NAPAA | 305774-339838 | ε-Poly-L-lysine | NRP | 100% | Antimicrobial |
| 2.3 | Terpene | 492890-513765 | Carotenoid | Terpene | 28% | Antioxidant |
| 4.1 | T3PKS | 53022-94104 | Saquayamycin A | Polyketide | 5% | Antimicrobial |

**Table S9.** Biosynthetic gene clusters in *Nesterenkonia aurantiaca* CMS1.6.

| Cluster ID | BGC type | Location (bp) | Most similar known cluster | Class | Similarity | Putative function |
|---|---|---|---|---|---|---|
| 3.1 | Terpene | 261588-282463 | Carotenoid | Terpene | 28% | Antioxidant |
| 5.1 | T3PKS | 75930-117012 | No hits | N/A | N/A | Unknown |
| 6.1 | Ectoine | 38903-49301 | Ectoine | Other | 50% | Osmoprotectant |
| 6.2 | NAPAA | 151878-185726 | ε-Poly-L-lysine | NRP | 100% | Antimicrobial |

**Table S10.** Biosynthetic gene clusters in *Arthrobacter* sp. KK5.5.

| Cluster ID | BGC type | Location (bp) | Most similar known cluster | Class | Similarity | Putative function |
|---|---|---|---|---|---|---|
| 4.1 | T3PKS | 55506-96801 | No hits | N/A | N/A | Unknown |
| 5.1 | NI-siderophore | 11655-42048 | Desferrioxamin B | Other | 50% | Metal chelation and transport |
| 6.1 | RiPP-like | 143257-154087 | No hits | N/A | N/A | Unknown |
| 8.1 | Ectoine | 80867-91271 | Ectoine | Other | 100% | Osmoprotectant |
| 10.1 | Terpene | 65697-86581 | Carotenoid | Terpene | 33% | Antioxidant |
| 14.1 | Betalactone | 23359-51531 | No hits | N/A | N/A | Unknown |
| 25.1 | Betalactone | 14021-33849 | Xantholipin | Polyketide | 4% | Antimicrobial |
| 29.1 | NAPAA | 1-21961 | Stenothricin | NRP, Cyclic depsipeptide | 31% | Antimicrobial |

**Table S11.** Biosynthetic gene clusters in *Aeromicrobium* sp. CF3.5.

| Cluster ID | BGC type | Location (bp) | Most similar known cluster | Class | Similarity | Putative function |
|---|---|---|---|---|---|---|
| 1.1 | NAPAA | 68663-102-490 | Stenothricin | NRP Cyclic depsipeptide | 27% | Antimicrobial |
| 1.2 | T3PKS | 362325-403395 | Alkylresorcinol | Polyketide | 100% | Antioxidant, antimicrobial |
| 1.3 | Redox cofactor | 524091-546826 | No hits | N/A | N/A | Cofactor |
| 2.1 | NI-siderophore | 543783-574125 | Desferrioxamine E | Other | 75% | Metal chelation and transport |
| 6.1 | Thioamitides | 59498-81971 | Kedarcidin | NRP+polyketide | 2% | Antimicrobial |
| 6.2 | T1PKS | 82717-128317 | Amycolamycin A/B | Polyketide | 12% | Antimicrobial |

**Table S12.** Biosynthetic gene clusters in *Aeromicrobium* sp. CF4.19.

| Cluster ID | BGC type | Location (bp) | Most similar known cluster | Class | Similarity | Putative function |
|---|---|---|---|---|---|---|
| 1.1 | NI-siderophore | 601562-631367 | FW0622 | Other | 50% | Metal chelation and transport |
